# Anemonefish have finer color discrimination in the ultraviolet

**DOI:** 10.1101/2022.12.01.518784

**Authors:** Laurie J. Mitchell, Amelia Phelan, Fabio Cortesi, N. Justin Marshall, Wen-sung Chung, Daniel C. Osorio, Karen L. Cheney

## Abstract

In many animals, ultraviolet (UV) vision guides navigation, foraging, and communication, but few studies have addressed the contribution of UV vision to color discrimination, or behaviorally assessed UV discrimination thresholds. Here, we tested UV-color vision in an anemonefish (*Amphiprion ocellaris*) using a novel five-channel (RGB-V-UV) LED display designed to test UV perception. We first determined that the maximal sensitivity of the *A. ocellaris* UV cone was at ∼386 nm using microspectrophotometry. Three additional cone spectral sensitivities had maxima at ∼497, 515, and ∼535 nm, which together informed the modelling of the fish’s color vision. Anemonefish behavioral discrimination thresholds for nine sets of colors were determined from their ability to distinguish a colored target pixel from grey distractor pixels of varying intensity. We found that *A. ocellaris* used all four cones to process color information and is therefore tetrachromatic, and fish were better at discriminating colors (i.e., color discrimination thresholds were lower, or more acute) when targets had UV chromatic contrast elicited by greater stimulation of the UV cone relative to other cone types. These findings imply that a UV component of color signals and cues improves their detectability, that likely increases the salience of anemonefish body patterns used in communication and the silhouette of zooplankton prey.

## Introduction

Many animals have ultraviolet (UV) sensitive (<400 nm) photoreceptors [reviewed by (1, 2)], and UV-vision contributes to behaviours including foraging (3, 4), celestial navigation (5, 6), mate selection (7–11), individual recognition (12), and aggressive displays (13, 14). But partly due to the technical challenge of producing UV-stimuli (15), few studies have tested its contribution to color discrimination, and it is unclear how UV sensitivity compares to that of visible range photoreceptors.

In vertebrates, cone photoreceptors in the retina mediate color vision. Most teleost fishes, reptiles, and birds have two morphological cone types: single cones and double cones. The latter is formed by the fusion of two cone cells (16–18). Photoreceptor spectral sensitivity is primarily determined by its photopigment(s) comprised of a G-coupled receptor opsin protein bound to a carotenoid-derived (Vitamin A1 or A2) chromophore, and can be modified by light-filtering in the eye (19, 20). Color vision can be defined as the ability to discriminate lights by their spectral composition regardless of their relative intensity. This requires a comparison of signals from different spectral types of photoreceptors, typically by chromatic opponent neurons (21, 22). To fully encode spectral information, an eye with n spectral receptor types require at least n-1 opponent mechanisms plus an achromatic (or luminance) mechanism (21). For example, birds are proposed to have UVS – SWS, SWS – MWS/MWS + LWS, and MWS – LWS (UVS, ultraviolet-sensitive; SWS, short-wavelength-sensitive; MWS, medium-wavelength-sensitive; and LWS, long-wavelength-sensitive) opponent systems (23, 24). The chromaticity (roughly hue and saturation) of a color can be defined by its location within an n-1 dimensional color space whose cardinal axes correspond to the activation of opponent signals (21), such as the 2-dimensional Maxwell’s triangle used for humans (25). Compared to human trichromatic color vision, adding a fourth cone allows tetrachromacy which expands the color gamut and can be represented by a tetrahedral color space (26, 27).

The presence of n-cone types does not mean that an animal has n-chromatic vision. For example, if the outputs of two spectral types of cones are summed, or a given cone type does not contribute to color discrimination. Thus, it has recently been found that the forward-looking retina of larval zebrafish (*Danio rerio*) forms a ‘strike zone’ (28). This zone contains a high density of enlarged UVS cones, which are suited to detecting zooplankton prey but make little contribution to chromatic opponency (28, 29). More generally, it is unknown whether UV chromatic contrast provides any major benefit to color discrimination in fishes, or in many animals for that matter, despite the presence of ample UV in many environments, including coral reefs, streams, grasslands, and forests (2, 30, 31).

Here we tested the UV and non-UV color vision capabilities of the false clown anemonefish, *Amphiprion ocellaris* (Figure 1A). Anemonefishes (genus, *Amphiprion*) are renowned for their symbiosis with sea anemones (Actiniaria spp.) (32) and their strict social hierarchy, which is determined by sex and body size (33, 34). Recently, anemonefishes were shown to have seven cone visual pigments, six of which are expressed in the adult *A. ocellaris* retina, where they produce four spectral types of cone (35). However, the exact cone spectral sensitivities in *A. ocellaris* were unknown, but estimated wavelengths of maximal sensitivity (λmax) according to their assigned opsin(s) indicate one type of single cone containing a mix of UV-sensitive/SWS1 (est. 368 – 370 nm λmax) and violet-sensitive/SWS2 (est. 406/407 nm λmax) opsins, and three types of double cones with blue-green RH2 opsins (RH2B est. 497 nm λmax and RH2A est. 516-523 nm λmax) in two separate members, and a third type with a mix of RH2A and LWS opsins (est. 560/561 nm λmax) (33).

**Fig. 1.**
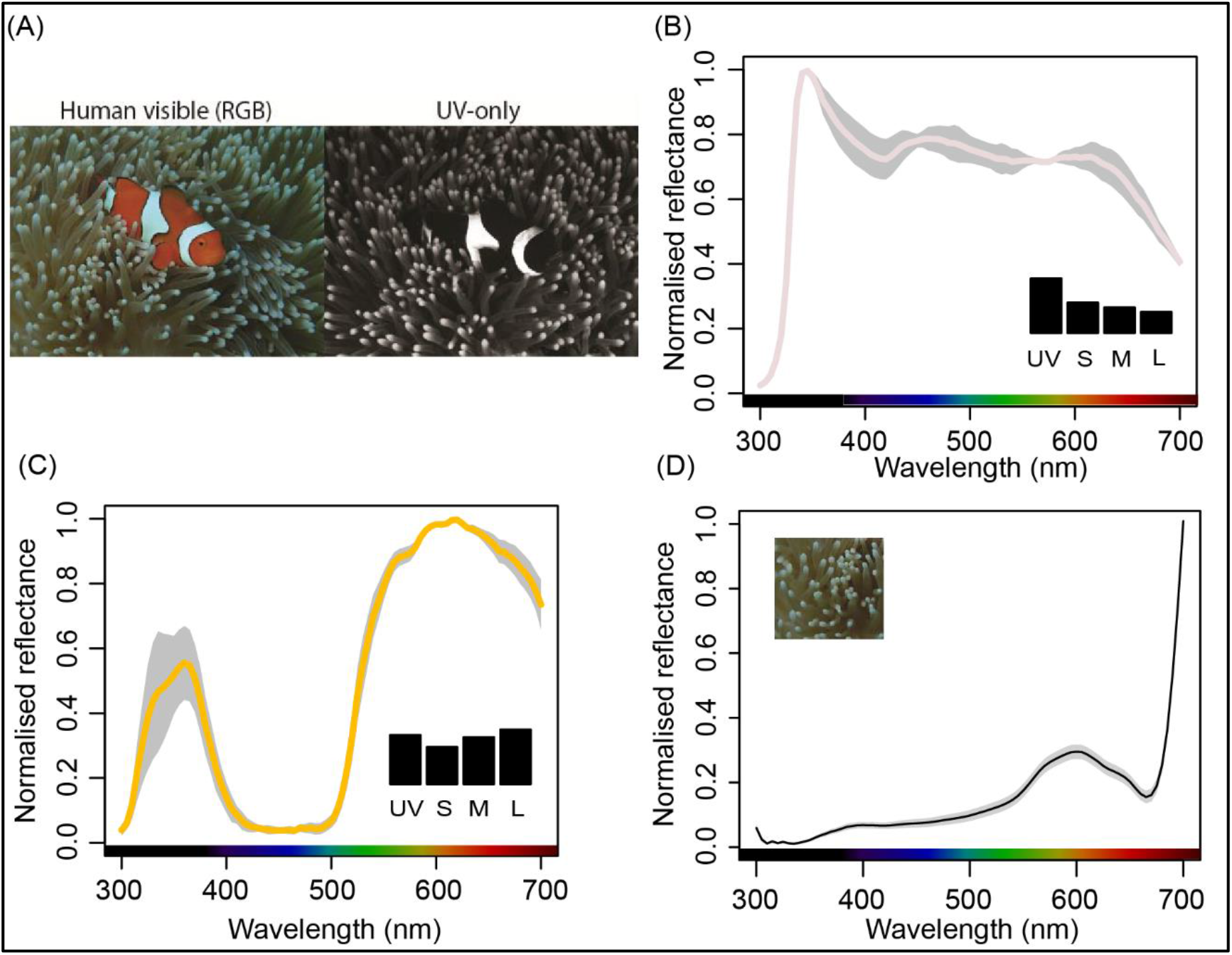
Colors of anemonefish skin and host sea anemone tentacles. (A) Underwater photographs of anemonefish taken in the human visible (RGB) spectrum and UV-only. Normalised spectral reflectance shown for *A. ocellaris* (B) UV-white bars, (C) UV-orange, and (D) sea anemone (*Stichodactyla gigantea*) tentacles. Lines represent mean spectral reflectance (n = 3), and shaded areas depict s.e.m. bounds. Note: images in (A) depict *A. percula*, the sister species of *A. ocellaris*, which shares an almost identical appearance and spectral reflectance.

The retinae of anemonefish are unusual from that of more commonly studied teleosts, such as goldfish (*Carassius auratus*) and zebrafish (both in the family, Cyprinidae), which have two single cones maximally sensitive to UV (SWS1 pigment, 355 – 365 nm λmax) and violet-blue (SWS2 pigment, 415 – 450 nm λmax), and double cones with separate ‘red’ (LWS) and ‘green’ (RH2) members (36, 37). As in larval zebrafish, anemonefish UV/violet cones are most abundant in a region of highest acuity or area centralis, but here, rather than being for prey detection, UV-sensitivity is thought to enhance the chromatic contrast of their UV-orange and UV-white colors for intraspecies communication (38) (Fig 1 A-C). Underwater UV-photography revealed that the light ‘white’ bars of *A. percula*, a sister species of *A. ocellaris*, have strong UV-contrast against adjacent dark ‘orange’ skin and the background of sea anemone tentacles (Fig 1A). Spectrometry of *A. ocellaris* skin confirms that both colors reflect UV: UV-white and UV-orange (Fig 1B and C). The natural background of anemonefishes is the tentacles of a host sea anemone (e.g., *Stichodactyla gigantea*) whose pronounced UV absorption below ∼350 nm (Fig 1D) coincides with the peak reflectance of anemonefish UV/white (Fig 1B), resulting in a dim appearance of the anemone compared to the fish’s bright UV/white bars (Fig 1A).

To investigate the colour discrimination capabilities of anemonefish to UV and non-UV signals, we first confirmed the sensitivity of all four cones using microphotospectrometry, and then conducted a behavioral experiment to examine how the three types of double cones, and the UV cones, contribute to color discrimination. Specifically, we asked whether discriminability was better for colors with higher UV chromatic contrast than those without (i.e., non-UV colors).

To do this, we used nine different sets of test colors produced by a five-channel (RGB-V-UV) LED display (15) customized to the *A. ocellaris* visual system. The contribution of photoreceptors to color vision were evaluated by the Receptor Noise Limited (RNL) model (19), which fits the psychophysical data of many species by assuming that color discrimination is set by chromatic opponent mechanisms whose performance is limited by noise arising in the photoreceptors (39, 40). The discrimination threshold (1ΔS) corresponds to a minimally distinguishable difference between two colors (19). Departures from RNL model predictions can provide evidence for post-receptoral mechanisms affecting color vision, including lateral inhibition (i.e., excited neurons suppressing neighboring neuron activity) (41), chromatic opponent and higher-level processes such as color categorization (42, 43).

## Results

### Spectral sensitivities of A. ocellaris

We first measured the lens transmittance and photopigment spectral sensitivities of *A. ocellaris*. The lens absorbed some UV wavelengths, with 50% transmission (T50) at 322, 340 and 341 nm in the three fish measured (mean T50 = 334 nm; Fig 2A). Microspectrophotometry (MSP) of the cone pigments found one type of single cone (U) with a maximal wavelength sensitivity (λmax) in the UV (mean λmax: 386 ± 5.0 nm; n = 4, N = 4 fish; Fig 2A). The three additional spectral cone types are double cones with λmax values of about 497 nm (M1), 515 nm (M2) and 531/538 nm (L) (Fig 2A). These photoreceptors could be assigned specific visual pigments according to their previously identified opsin protein component (Fig 2A) (35). One type of rod photoreceptor was present (mean λmax value = 502 ± 4.0 nm; n = 7 cells, N = 4 fish; Supplementary Figure 1). All double cone absorbance spectra fitted a retinal (Vitamin A1) derived chromophore visual pigment template, while the single cone absorbance was likely due to the coexpression of UV-and violet-sensitive visual pigments, as has previously been shown to be the case in *A. ocellaris* (35). In vivo photoreceptor spectral absorbance curves were given by the product of lens transmission and photoreceptor spectral absorbance measurements (Fig 2A). Two measurements hinted at a third MWS (M3) double cone type (508/509 nm λmax, N = 1 fish; Supplementary Figure 1), but the spectral overlap with the M2 cone meant it is unlikely that M3 makes a separate contribution to color vision. Discrimination threshold estimates for target sets that included its input or substituted it with M2 provided a worse overall fit (see Supplementary Figure 2).

**Fig. 2.**
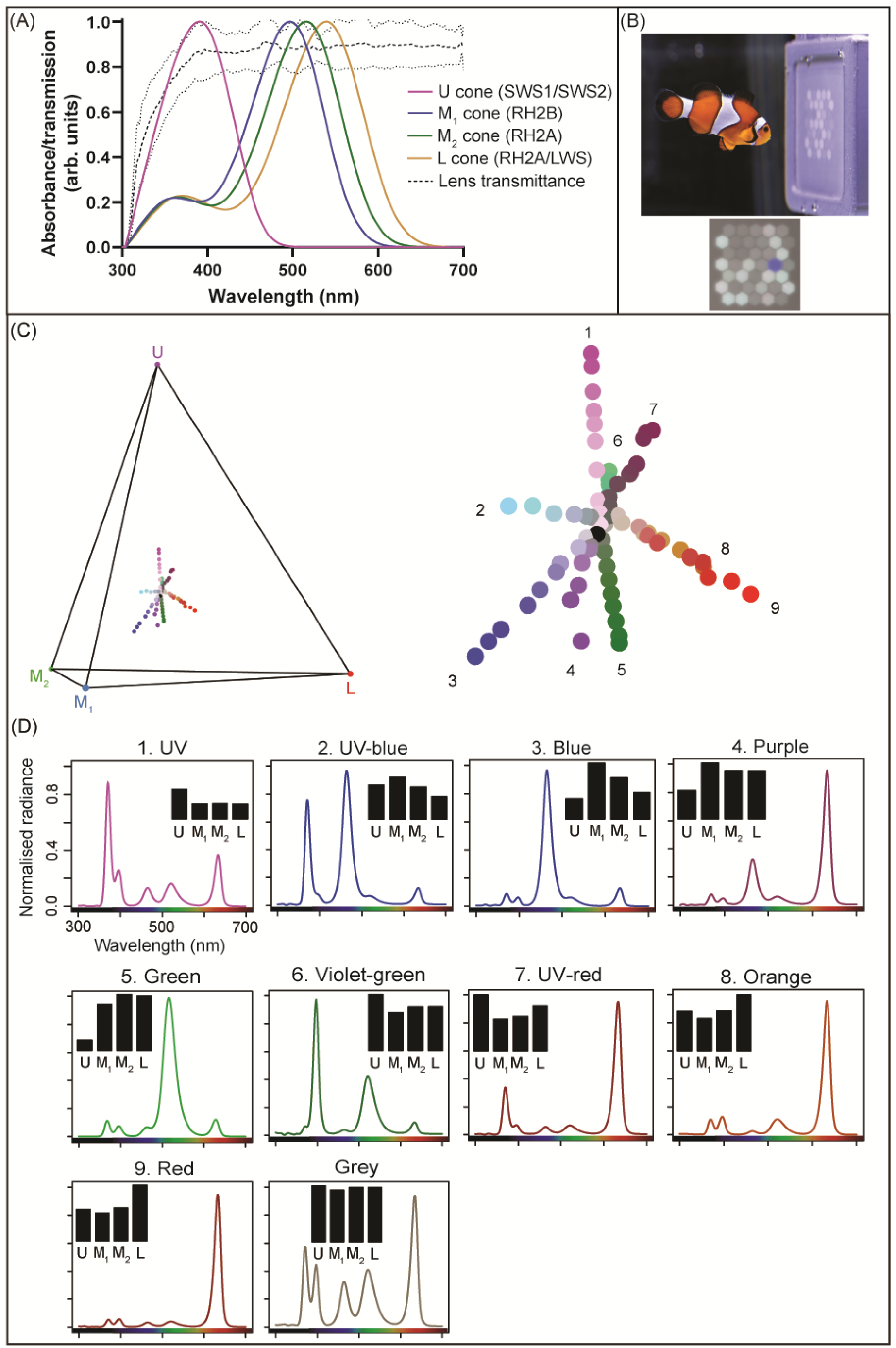
Cone spectral sensitivities of *A. ocellaris* and the LED display used for testing the discriminability of nine different sets of colors. (A) Normalised average lens transmission of *A. ocellaris* (N=3), and absorbance curves (in arbitrary units) fitted to the averaged absorbance spectra of the cones (N=9). In parentheses are the identities of (co-)expressed visual opsins that principally determine the spectral sensitivities of *A. ocellaris* cones (35): ‘SWS1/SWS2’ = UV/short-wavelength-sensitive 1 and violet/short-wavelength-sensitive 2 (U) (λmax: 386 ± 5.0 nm, n = 4, N = 4 fish), ‘RH2B’ = medium-wavelength-sensitive rhodopsin-like 2B (M1) (λmax: 496.7 ± 3.0 nm, n = 3, N = 3 fish), ‘RH2A’ = medium-wavelength-sensitive rhodopsin-like 2A (M2) (λmax: 515 ± 2.1 nm, n = 6, N = 3 fish), and ‘RH2A/LWS’ = long-wavelength-sensitive (L) (λmax: 531/538 nm, n = 2, N= 1 fish). Thin broken lines depict the upper and lower bounds of the standard deviation for lens transmittance. Supplementary Figure 1 gives individual absorbance spectra for all photoreceptors. (B) RGB-V-UV LED display with an example stimulus. Fish were trained to discriminate a target dot from distractor dots and peck it to receive a food reward. (C) Tetrahedral receptor space of *A. ocellaris* showing the nine tested color sets (and achromatic point in black), and (D) the normalized spectral radiance of nominally saturated example targets and the average grey distractor. Bar plots depict the relative receptor stimulation (quantum catches) evoked by colors or grey for *A. ocellaris* against the PTFE screen of the LED display. ‘U’ = ultraviolet-sensitive, ‘M1’ = medium-wavelength-sensitive 1, ‘M2’ = medium-wavelength-sensitive 2, and ‘L’ = long-wavelength-sensitive cone types. Note, color names have no implication for their appearance to the fish. Image credit: *A. ocellaris* taken by Valerio Tettamanti.

### Color discrimination thresholds

Anemonefish were trained to peck a rewarded target pixel that differed in chromaticity from grey distractor pixels (Fig 2B) (*44, 45*). The grey level of the distractor pixels was varied to prevent the use of brightness information. We then measured the fishes’ accuracy for nine sets of target colors (Fig 2C, D). Each set lay on a line radiating from the central achromatic (grey) point in the anemonefish color tetrahedron, so that they varied in saturation but not hue. Four of the nine colors we collectively refer to by ‘UV colors’ (UV, UV-blue, UV-red, and Violet-green; Fig 2C) had increasing UV/violet LED emission, while the five remaining (Blue, Green, Red, Orange, and Purple; Fig 2C) were ‘non-UV colors’ without increasing UV saturation. Four of the test colors also excited spectrally non-adjacent receptors more than intermediate receptors (Violet-green, UV-Red, Orange, and Red), and so were non-spectral colors, i.e., the human equivalent of ‘purple’ (e.g., [27]), which could not be matched by a mixture of a monochromatic light with grey (Fig 1D).

We conducted a total of 3921 test trials (N = 11 fish, n = 9 color sets, N = 84 colors, n = 8 – 17 trials per target, mean = 10 trials). Our (50%) probability of correct choices for determining the threshold is more accurate than more commonly used (70% to 75%) in paired choice tests (*46*), as the experiment involved a choice of 1 out of 38 pixels. Color differences were specified by the RNL model for anemonefish (where 1 Δ*S* is defined as the discrimination threshold). Because noise levels in *A. ocellaris* cones are unknown, we found the best fit of the RNL model to the data (*47, 48*). This best fit predicted receptor noise of σ = 0.11 in the UV (single) cones, and σ = 0.14 for the three types of double cone, which gave the following thresholds for the nine test colors: blue (mean ± s.e.m = 1.5 ± 0.1 Δ*S*), purple (1.6 ± 0.07 ΔS), green (1.2 ± 0.1 ΔS), red (1.0 ± 0.04 ΔS), orange (0.8 ± 0.07 ΔS), UV-blue (0.8 ± 0.07 ΔS), UV (0.8 ± 0.09 ΔS), Violet-green (0.4 ± 0.02 ΔS), and UV-red (0.9 ± 0.05 ΔS). These values compare to a receptor noise estimate of σ = 0.05 in another reef fish [43, 45, 49]. Because vectors are origin bound, the vectors representing colors in anemonefish color space at 180° with each other can be considered as complementary pairs, and when mixed in equal proportions, should be achromatic (*50*). Based on the approximately polar azimuth angles among some color thresholds (φ ± 180; Fig 3B, C), we were able to identify two pairs of what are likely complementary colors: *1*) Green (φ = 297°) and UV (φ = 127), and *2*) Purple (φ = 128) and Violet-green (φ = 298).

**Fig. 3.**
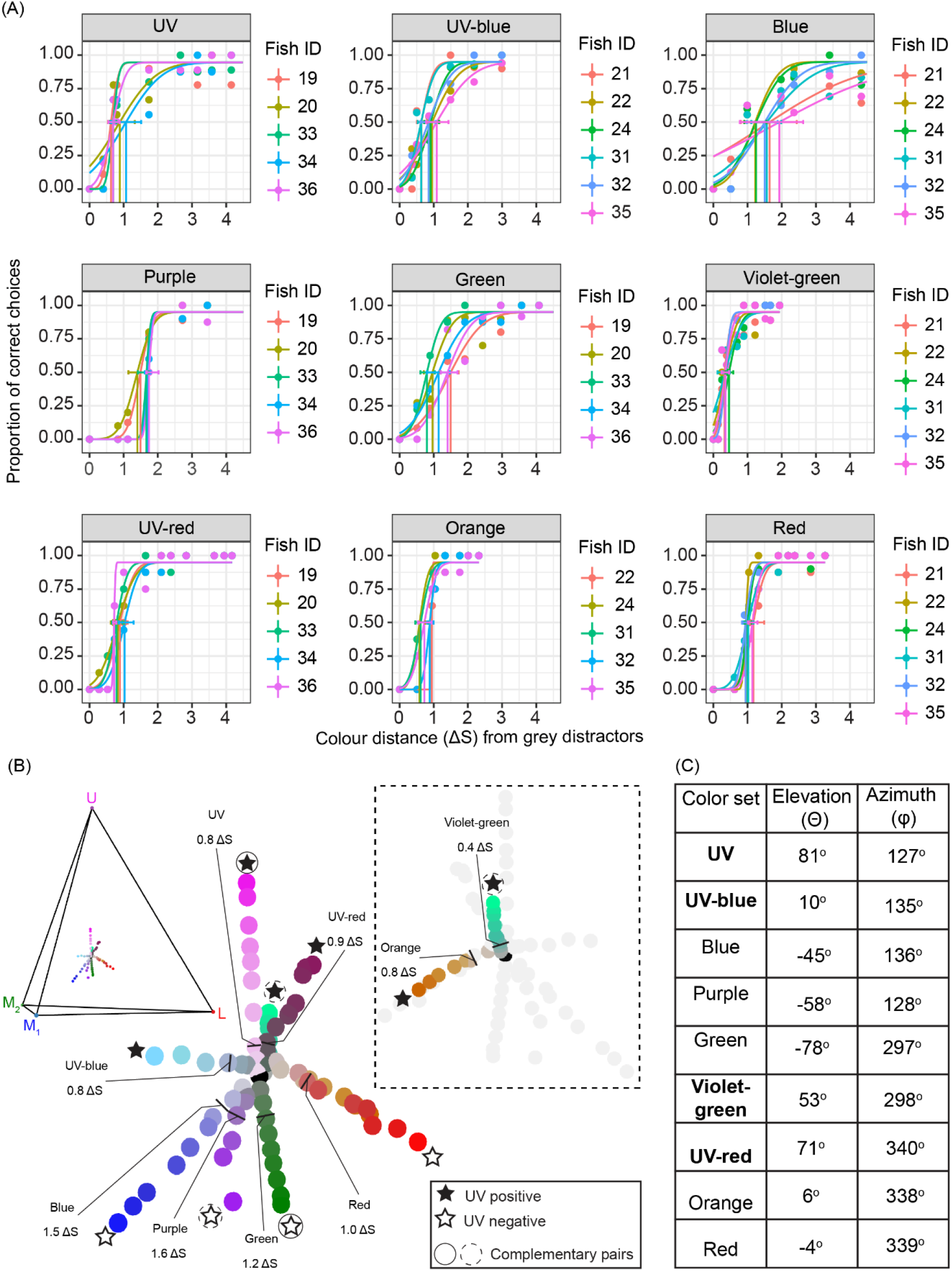
Anemonefish color discrimination thresholds and their hue angles in tetrahedral space. (A) Color discrimination thresholds shown as a function of the proportion of correct choices by anemonefish for targets with a range of chromatic contrasts (Δ*S*). Error bars denote 0.95 CIs. Discrimination thresholds are values calculated per fish (Fish ID) and are demarked by vertical lines. Each plotted point represents the mean proportion of correct choices from one fish (n = 8 to 17 trials). Note, the x-axis has been truncated to ≤4.5 Δ*S* for presentation purposes. For curves showing all the data see Supplementary Figure 3. (B) Relative positions of equal discriminability marked by intersecting lines in the receptor space of *A. ocellaris* for target sets of colors that varied in chromaticity (Δ*S*) in four principal directions representing the relative stimulation of the ultraviolet-(U), medium 1-(M_1_), medium 2-wavelength sensitive (M_2_), and long-wavelength sensitive (L) photoreceptors. Star symbols and circles adjacent to color sets indicate whether UV receptor input was needed for discriminating threshold colors from grey as determined by a positive elevation vector angle (Θ >0), and which colors potentially form complementary pairs according to polar azimuth vector angles (approximately φ ± 180°) respectively. Intersecting lines and adjacent numbers refer to the average discrimination threshold of each target set. The black dot represents the average grey distractor spectra which the receptor space is centered on (i.e., the origin). The inset is the reverse view of the receptor space to show the Orange and Violet-green sets clearly. Δ*S* was calculated using a receptor noise standard deviation (σ) of 0.11 for single cones and 0.14 for double cones. (C) Summary of hue angles calculated in degrees using *XYZ* Cartesian coordinates according to noise-corrected threshold ΔS values. Elevation ‘Θ’ describes the ‘vertical’ angle (−90° ≤ Θ ≤ 90°) of threshold vector position above (positive) or below (negative) the plane of chromaticity facilitated by the double cones (M_1_, M_2_, L), where a 90° direction corresponds to the U cone vertex. Azimuth is the ‘horizontal’ angle (0° ≤ φ ≤ 360°) from the L cone at 0°/360° with the U cone axis normal to the equatorial plane. See Methods for hue angle equations and Supplementary Table 1 for relative quantum catches of cones and *XYZ* Cartesian coordinates for vectors. UV colors (i.e., those produced by higher UV/violet LED emission) are shown in bold.

To examine the possibility that only a subset of cones contributes to color vision in anemonefish and one cone type might only serve achromatic tasks (i.e., trichromacy), we also compared the tetrachromatic model fit to that of four possible trichromatic models, where the input from one cone type was systematically dropped. None of the trichromatic models predicted the discrimination of all test colors from grey to an expected 1 ΔS (two-way ANOVA, range of estimated mean difference = 0.101 to 0.40, F = 18.1, all *p* < 0.05; Fig 4A). The tetrachromatic model (UM_1_M_2_L) had the closest fit (two-way ANOVA, range of estimated mean difference = −0.006, F = 18.1, *p* = 0.910; Fig 4A), which was followed by the UM_1_L model missing the M_2_ cone (two-way ANOVA, estimated mean difference = 0.101, F = 18.1, *p* = 0.046; Fig 4A).

**Fig. 4.**
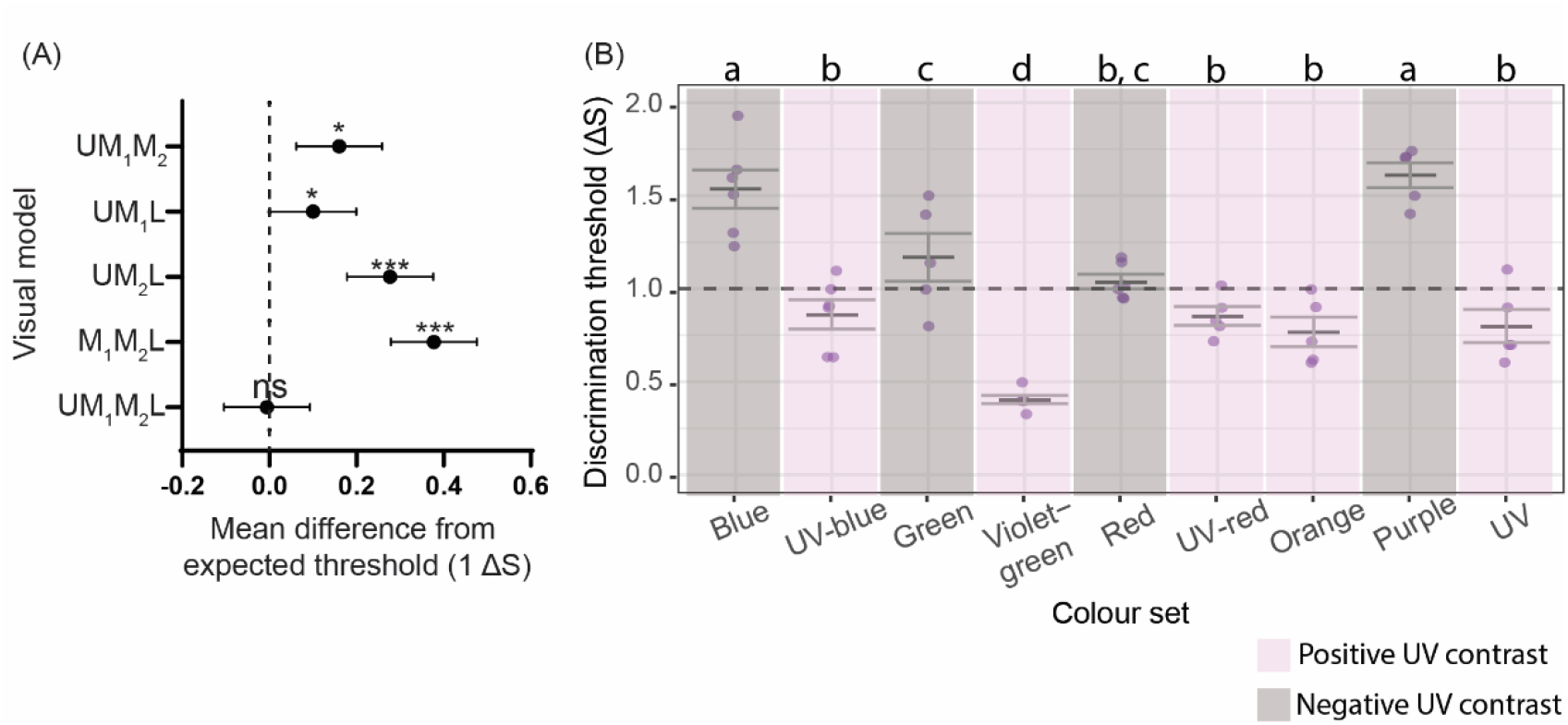
Discrimination threshold fit comparisons among different RNL model scenarios and color sets. (A) Comparison between tetrachromatic (UM_1_M_2_L) and trichromatic modelling in the accuracy of predicting color discrimination thresholds. For each model, the individual thresholds (ΔS) per color set were subtracted from the expected threshold value of 1 ΔS (demarked by the broken line), and the overall average difference was calculated. Visual models either included input from all cones (tetrachromatic) or dropped input from one of the four cone types (trichromatic): UV-sensitive cone ‘U’, medium-wavelength sensitive cone 1 ‘M_1_’, medium-wavelength sensitive cone 2 ‘M_2_’, or long-wavelength sensitive cone ‘L’. Error bars indicate upper and lower 0.95 confidence limits. ‘ns’ indicates *p* > 0.05, ‘*’ indicates *p* = 0.05 > 0.001, and ‘**’ indicates *p* <0.001. All *p*-values were calculated by two-way ANOVA with Fisher’s Least Significant Difference test. See Supplementary Figure 4 for a multiple model comparison of individual color discrimination thresholds. (B) Pairwise comparisons of *A. ocellaris* discrimination thresholds (ΔS) between color sets with or without UV contrast. Letters denote statistical significance (*p* < 0.05) between groups. Grey bars indicate mean values while error bars represent ± 1 s.e.m. The broken, horizontal line demarks the RNL model assumed threshold (ΔS = 1). ΔS were calculated using a receptor noise standard deviation (σ) of 0.11 for single cones and 0.14 for double cones. Statistical significance was calculated using a Linear Mixed Effects Model (LMM) with multiple paired comparisons. See Supplementary Data 1for full LMM results.

### Lower discrimination thresholds induced by UV

Colors with positive UV chromatic contrast were identifiable by a positive elevation angle (Θ > 0) and had significantly lower thresholds (UV = 0.8 ΔS, Θ = 81°; Violet-green 0.4 ΔS, Θ = 53°; UV-red 0.9 ΔS, Θ = 71° and UV-blue 0.8 ΔS, Θ = 10°; Fig 3B and 4B) than three of the five with negative UV chromatic contrast (Θ < 0) (Blue 1.5 ΔS, Θ = −45°; Green 1.2 ΔS, Θ = −78°, and Purple 1.6 ΔS, Θ = −58°; Fig 3B and 4B) (LMM, all paired comparisons *p* < 0.05; Fig 4B). Red (1.0 ΔS, Θ = −4°), Orange (0.8 ΔS, Θ = 6°), and UV-blue thresholds had low UV chromatic contrast and were located near the equatorial plane (Θ = 180° ± 10.0°) (Fig 3B). Both identified pairs of complementary colors had positive and negative UV chromatic contrasts, where the former had significantly lower discrimination thresholds for UV/Green (LMM, estimate ΔS_UV – Green_ = −0.40, se = 0.11, z = −3.22, *p* = 0.012; Fig 4B) and Violet-green/Purple (LMM, estimate ΔS_Violet-green – Purple_ = −1.21, se = 0.11, z = −11.0, *p* <0.0001; Fig 4B).

Psychometric functions of the nine color sets differed significantly, with blue, purple, green, UV, and UV-blue sets having more gradual functions (Fig 3A) than orange, red, violet-green, and UV-red sets (binomial generalized linear mixed-effects model/GLMM, all *p* ≤ 0.01; for GLMM results and pairwise comparisons see Supplementary Data 1). A more gradual incline was indicative of a higher error rate for relatively high Δ*S* targets, and a higher Δ*S* asymptote for discrimination performance. The differences in these functions were not attributable to the order in which colors were presented and followed no obvious pattern. Because some color sets were introduced later than others in the experiment (refer to Methods for the introduced order of color sets), this may have contributed to discrepancies in psychometric functions and thresholds. To determine whether the presentation-order of color sets affected discrimination performance, we reassessed two fish per color set at the end of the experiment. We found that all color sets had either no change or only minor differences in discrimination thresholds (range = 0 – 0.4 Δ*S* shift, mean ± s.e.m. = 0.1 ± 0.02 Δ*S* shift; see Supplementary Figure 5 for psychometric curve comparisons).

### Decision latency

The modelled detectability of the test color against the background (Δ*S*) strongly predicted choice latency for all color sets except orange (LMM, all *p* < 0.001; Supplementary Data 1) but no significant differences were detected in the latency between colors.

## Discussion

We have shown that the anemonefish UV cone which has a peak sensitivity (λ_max_) at 386nm contributes to tetrachromatic color vision with the three spectral types of double cones (λ_max_ 495nm, 515nm and c. 535nm). This suggests that the cones containing all four combinations of the main pigment types (SWS1/SWS2, RH2B, RH2A, RH2A/LWS) (*35*) contribute separately to color vision, and that the UV cone has a comparatively high sensitivity. These conclusions are striking given the evidence for co-expression of the RH2A and LWS pigments in the same double cone member, and of the SWS1 and SWS2 pigments in the single cones (*35*). There is experimental evidence for tetrachromacy in a few other species, including goldfish (*51, 52*) and chicken (*53*), but to our knowledge this is the first demonstration by testing the minimally saturated hues that can be distinguished from grey, which clearly suggests that anemonefish have a 3-D color space. This advance was made possible by our five channel LED display customized to anemonefish vision (Fig 2B). The display also allowed us to show that anemonefish can discriminate a wide variety of non-spectral colors from grey, which would be highly difficult with monochromatic test lights (*15, 27*).

A major finding was the importance of the UV receptor in anemonefish color vision. By providing a rare comparison of discrimination thresholds among relatively unexplored UV-regions of animal color space, we show that positive UV chromatic contrast improved the discrimination of four of the nine color sets (UV, UV-blue, Violet-green, and UV-red). The estimated noise (σ = 0.11) in the UV cone was lower than in the three double cones (σ = 0.14). Furthermore, fitting the RNL model revealed that the discrimination thresholds for these colors were substantially lower than three of the nine color sets (Blue, Green, and Purple) which had negative UV chromatic contrast, i.e., the relative stimulation of the single cone was lower than all three double cones. This difference in discriminability was most convincingly shown by the complementary color pairs UV/Green and Violet-green/Purple, which are theoretically equidistant in ΔS from grey (*50*) but had large disparities in psychophysical threshold distances. This asymmetry between color discrimination thresholds cannot be directly attributed to noise in the early stages of the visual pathway such as photoreceptor noise or chromatic opponent neurons in the retina. One possibility is that activation of the UV receptor suppresses noise in the visual pathway or enhances the saliency of colors for anemonefish. The high sensitivity to violet-green, which was found in all six of the fish tested is consistent with the heightened saliency of this color.

### Positive UV chromatic contrast benefits anemonefish color discrimination

UV-contrast sensitivity has been reported in multiple animals, such as common goldfish (*41, 52*), zebrafish (*54*), budgerigars (*Melopsittacus undulatus*) (*24*), and hummingbirds (*Selasphorus platycercus*) (*27*). Previously, *A. ocellaris* demonstrated an ability to detect UV targets (*15*), which did not strictly require color vision, and like the larval zebrafish (*28*), one anemonefish (*Amphiprion akindynos*) has been found to have UV cones most concentrated in the frontal visual field (i.e., the centrotemporal retina) (*38*). It is unknown whether *A. ocellaris* also has a peak UV cone abundance in its centrotemporal retina; however, this seems quite likely given that they share multiple key features with their larger cousin *A. akindynos* (*38*), including similar cone spectral sensitivities and photopigment diversity (*35*), and common ecological aspects (life history, sea anemone habitat, social hierarchy, diet). In the zebrafish, UV cones seem to make little input to color vision (*28*), while in anemonefish UV cones make a disproportionately strong input. This strong UV contribution to color discrimination suggests that anemonefish color patterns are highly salient to conspecifics and could benefit UV-signaling used in social communication (*55*).

### Color preferences and ecological significance of UV

An innate (or learnt) UV-preference in *A. ocellaris* could explain their acute discrimination, where a higher attention to UV may be influenced by the color of their food or conspecifics. The white bars of *A. ocellaris* appear to have strong UV contrast against adjacent dark orange skin and sea anemone tentacles, as was shown to be the case in *A. akindynos* (*38*). Indeed, juvenile anemonefish (*A. akindynos*) have a distinct UV coloration shown to signal subordinance (*55*).

Other suggested functions of anemonefish color patterns include warning coloration (*56*), camouflage (*56*), species recognition (*57*), and mate recognition (*58*). Future studies on the function of anemonefish coloration should include the UV and not restrict their spectral analysis to longer wavelengths in the human visible spectrum. Moreover, attention should be paid to how the appearance of these colors change with increasingly common sea anemone bleaching events and the implications this may have for signaling efficacy. Another potential basis for a UV preference in anemonefish could be to detect the UV-contrast of their common prey (zooplankton) which can either scatter or absorb UV (*28, 59, 60*). Larval anemonefish (*A. biaculeatus*) can solely rely on UV illumination (peaking at 365 nm) for detecting prey (*61*), which might be attributed to an achromatic UV channel as in larval zebrafish (*28*). More generally, highly sensitive UV vision could help maintain the detectability of UV signals in habitats with reduced UV photon availability (e.g., deep water, dense foliage cover, heavy overcast), as suggested in goldfish (*62*). Anemonefishes typically inhabit shallow coral reefs (about 1 – 15 meters) where UV is abundant; however, a similar mechanism for facilitating acute UV discrimination might exist in other marine fishes and benefit UV vision in deeper habitats even beyond 100 meters and at a maximum of 200 meters, where a conservative estimate indicates enough UV photons could sustain the visual sensitivity of surface-dwelling fishes (*63*).

An aversion towards blue and purple might explain their high discrimination thresholds (∼1.5 Δ*S*) and variable psychometric functions. This variation may indicate individual differences in learning (e.g., categorical perception), or differences in attentiveness and motivation. Guppy (*Poecilia reticulata*) also poorly discriminate purple due to possible neophobia (*64*), and triggerfish have an aversion towards blue (*65*), which was explained based on its common use as an aposematic color, signaling unpalatability in reef invertebrates such as nudibranchs (*66*) and blue-ringed octopus (*67*).

## Conclusion

We found the discriminability of colors by anemonefish varied depending on UV contrast, where positive UV chromatic contrast had lower discrimination thresholds. Thus, it appears that UV vision can aid the detectability of fine-scale differences in chromaticity and might benefit the viewing of natural UV-reflective objects (e.g., colors of prey and conspecifics). This raises new questions regarding the identity and function of neural mechanism(s) in the retina and/or brain that facilitate highly sensitive UV color discrimination. Although we did not explicitly test the extent of cone opponency in anemonefish, their psychophysical thresholds were best explained when all four cone types contributed to color vision suggesting tetrachromacy. The enhancement of UV color saliency in anemonefish likely has an ecological benefit for communicative signaling using their UV colors.

## Materials and methods

### Animals and ethics statement

Anemonefish (*A. ocellaris*) (N = 20) were acquired from a local aquarium store (supplier Gallery Aquatica, Wynnum, 4178 QLD, Australia). We used N = 9 (female = 3, mean total length = 4.5 ± 0.5 cm; male = 6, mean total length = 3.5 ± 0.5 cm) for taking measurements of cone spectral sensitivities, and N = 11 (female = 11, mean total length = 4.9 ± 0.3 cm) for behavioral experiments. Fish were housed individually in recirculating aquaria (60×30×30 cm) at the Queensland Brain Institute at The University of Queensland, Australia, and all experiments were conducted in accordance with The University of Queensland’s Animal Ethics Committee guidelines under approval numbers QBI/304/16 and SBS/077/17. For anatomical measurements anemonefish were euthanized by immersion in MS222 (500 mg/L) for 10 minutes and subsequent decapitation.

### Underwater photography and measuring skin spectral reflectance

Underwater photographs of anemonefish (*Amphiprion percula*) were taken at Horseshoe Reef near Lizard Island on the Great Barrier Reef (Figure 1A). Photographs in the visible (RGB) were captured using an Olympus (TG-5) camera (with Nikon 60 mm Micro), while a UV-converted Nikon (D810) fitted with both a short-pass filter (UG 11) and far-red filter took UV images.

Reflectance spectra of *A. ocellaris* white and adjacent orange skin patches surrounding the head/operculum region were measured for three wild-caught, captive males (n = 2) and a female (n = 1), for which no major differences were found. Three replicate measurements per skin area (within roughly 0.5 cm^2^ – 1 cm^2^) were taken across a 300 – 700nm range using a spectrometer (USB4000 Ocean Optics) with a pulsed Xenon lamp (PX-2 Ocean Optics), and a bifurcated 200µm fibre optic cable (Ocean Optics). Reflectance measurements were captured out-of-water at a 45° angle directly against skin and relative to a Spectralon 99% diffuse reflectance standard (Lab-sphere, North Sutton, NH, USA). During the measuring process fish were restrained in a sea water-soaked towel to protect their eyes from the bright light and skin from desiccation. Exposure out of water was limited to a maximum of 1-minute, and post-measurement all fish recovered and were returned to their aquarium. Sea anemone reflectance was based on averaged (n = 10) measurements recorded in-situ using a submersible spectrometer (USB2000 Ocean Optics) with a 100µm fibre and relative to a 99% Spectralon reflectance standard positioned next to the anemone. Reflectance measurements of tentacles used natural daylight as a light source at midday during non-overcast conditions at ∼3m depth.

### Lens transmission of A. ocellaris

For the measurement of lens transmission in *A. ocellaris*, the lenses (n = 3 fish) were isolated from the hemisected eyecup and rinsed in PBS to remove any blood and vitreous. Spectral transmission (300-800 nm) was measured by mounting the lens on a drilled (1.0 mm diameter hole) metal plate between two fibers (50, 100 µm diameters) connected to an Ocean Optics USB4000 spectrometer and a pulsed PX2 xenon light source (Ocean Optics, USA). Light spectra were normalized to the peak transmission value at 700 nm (*68*), and lens transmission values were taken at the wavelength at which 50% of the maximal transmittance (T_50_) was attained (*68, 69*). No pigmented ocular media was observed.

### Photoreceptor spectral sensitivities of A. ocellaris

The spectral absorbance of *A. ocellaris* photoreceptors were measured using single-beam wavelength scanning microspectrophotometry (MSP). This procedure followed that outlined in detail elsewhere [see (*70, 71*)]. In summary, small pieces (∼1 mm^2^) of tissue were excised from the eyes of two-hour dark-adapted fish, then immersed in a drop of 6% sucrose (1X) PBS solution and viewed on a cover slide (sealed with a coverslip) under a dissection microscope fitted with an infra-red (IR) image converter. A dark scan was first taken to control for inherent dark noise of the machine and a baseline scan measured light transmission in a vacant space free of retinal tissue. Pre-bleach absorbance measurements were then taken by aligning the outer segment of a photoreceptor with the path of an IR measuring beam that scanned light transmittance over a wavelength range of 300-800 nm. Post-bleach scans were then taken after exposing the photoreceptor to bright white light for 60 seconds, and then compared to pre-bleach scans to confirm the presence of a labile visual pigment. Confirmed visual pigment spectral absorbance data was then analyzed using least squares regression that fitted absorbance data between 30% and 70% of the normalized maximum absorbance at wavelengths which fell on the long-wavelength limb. The wavelength at 50% absorbance was then used to estimate the maximum absorbance (λ_max_) value of the visual pigment by fitting bovine rhodopsin as a visual pigment template (*72, 73*). This absorbance curve fitting was performed in a custom (Microsoft Excel) spreadsheet, where the quality of fit of absorbance spectra between A1-and A2-based visual pigment templates was also visually compared. Individual scans were binned on their grouping of similar (≤10 nm difference) λ_max_ values, and then averaged and reanalyzed across fish to create mean absorbance spectra (Supplementary Data 3).

### LED display and stimuli calibration

To display the visual stimuli in our behavioral experiments we used a five-channel RGB-V-UV LED display [for full design details, see (*15*)]. Note, that the violet channel had an emission that emitted into the UV and violet, where it had higher overlap with the absorption curve of the UV cone but is referred to as ‘violet’ to distinguish it by name from the shorter wavelength ‘UV’ LED. The display itself was held within a waterproof, 3D-printed case, with a PTFE screen that acted as a light diffuser. A wide gamut of colors could be produced by modulating the relative outputs of each LED to color mix the different channels.

Target and distractor colors were chosen to test anemonefish color discrimination along nine different sets of chromatic contrast including: UV, UV-blue, blue, purple, green, violet-green, UV-red, orange, and red. We first measured the spectral radiance (µM cm^-2^ s^-1^ nm^-1^) of pixel colors using a spectrometer (Ocean Optics USB4000) with a 200 µm diameter UV-VIS fiber calibrated against a deuterium-halogen lamp (Mikropak DH2000-DUV, calibrated by Ocean Optics). An RPA-SMA holder (Thorlabs) maintained the fiber 1 mm directly above a pixel at a 90° angle.

The stimulus used for measuring discrimination thresholds was inspired by the Ishihara test of color vision deficiency (*44*), as per (*45*). Anemonefish were trained to discriminate a target pixel which differed in chromaticity from distractors (*43*). We ran the LED display via a Python script that pseudo-randomly assigned a target color to one out of 38-pixel coordinates, while the 37 remaining pixels were assigned as grey distractors. Note, we did not utilize the full-sized display due to fish being afraid of the LED display and not trainable when using a larger stimulus.

### Color selection and stimuli design

To estimate anemonefish photoreceptor excitation for target and distractor colors, receptor quantum catches ‘*q*’, were first calculated for each stimulus, ‘*S*’ (i.e., target and distractor radiance spectra in µM/cm^-2^/s^-1^/nm) viewed under well-lit conditions given by:

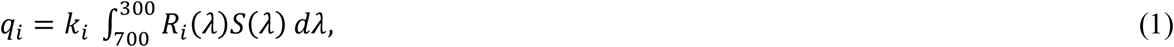

where *k* is a scaling coefficient for receptor adaption to the background ambient light, *S*_*b*_:

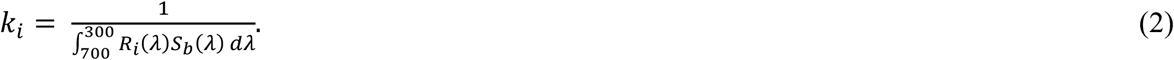

*R*_i_(*λ*) was the normalized spectral absorbance of a given receptor type ‘*i*’ (*i* = U, M_1_, M_2_, L) multiplied by lens transmittance, and ‘*λ*’ denoted wavelength (nm). *S*_*b*_(*λ*) was the spectral radiance of the PTFE display screen (between the pixels) with all LEDs turned off and measured from 5.0 cm in the experimental tank (for background radiance spectra see Supplementary Figure 6). This approach allowed for modelling spectral emission (from LEDs) rather than more commonly calculated for reflectance, as per (*74*). Integration was performed across the visible spectrum (i.e., 300 – 700 nm for *A. ocellaris*). Relative cone quantum catches (see Supplementary Data 2 for absolute quantum catches) were used to plot color loci in a tetrahedral color space (*26, 75*).

Next, we calculated the chromatic contrast or color distances (Δ*S*) of target colors relative to the average distracter spectra using the log receptor noise limited (RNL) model (*19, 46*). This provided a visual representation of how target and distractor colors appeared to *A. ocellaris*. The RNL model assumes: *1*) that 1 Δ*S* equates to a psychophysical threshold of one just noticeable difference between two stimuli of a given contrast, *2*) color vision is conveyed by chromatic mechanisms independent of achromatic visual processes, and *3*) that Δ*S* is determined by the differences in receptor stimulation (Δ*q*_*i*_) elicited by two viewed stimuli, that is only constrained by receptor noise levels (*e*_*i*_) for each of the four cone classes (*i* = 1, 2, 3, 4 for the U, M_1_, M_2_, L cone classes), or alternatively, for three cone classes in trichromat models (i = 1, 2, 3 for the U/M_1_/M_2_/L cone classes).

The contrast (Δ*q*_*i*_) for each receptor channel was calculated by,

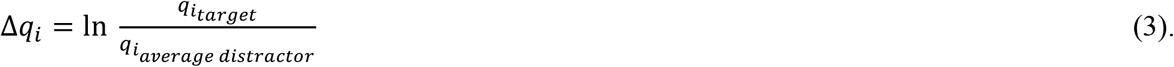

In the absence of direct noise measurements for *A. ocellaris* cones, we estimated cone noise levels (*e*_*i*_) by,

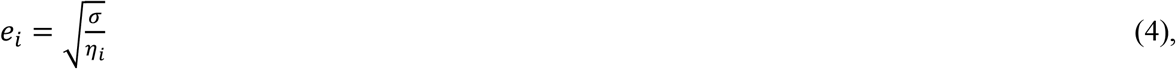

where ‘σ’, the numerator of the Weber fraction in (*48*), or ‘*v*_*i*_’ in (*19*) is the standard deviation of noise within a photoreceptor, and ‘η’ is the ratio of the given cone type. We initially assumed that cone noise levels in *A. ocellaris* were like those reported in triggerfish and used a σ-value = 0.05 (*49, 65, 76*). Based on the regular mosaic of one single cone surrounded by four double cones in the *A. ocellaris* retina (*35*), we used a relative cone abundance ratio of 1 : 2 : 1 : 1 (U : M_1_ : M_2_ : L) for a tetrachromatic visual system and 1 : 2 : 2 for a trichromatic visual system. The double cone ratio for the tetrachromatic scenario was based on in-situ hybridization experiments showing that *A. ocellaris* in our aquarium system express *RH2B* (M_1_) in one double cone member and either *RH2A* (M_2_) or *RH2A* with *LWS* (L) in the second double cone member (*38*).

Δ*S* in tetrachromatic visual space was calculated by:

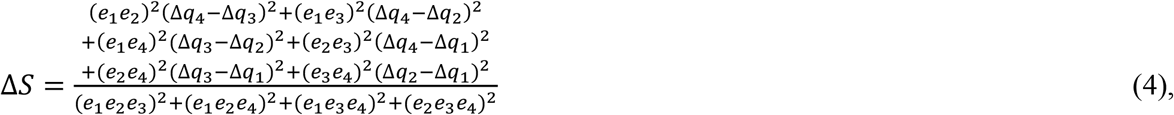

and in trichromatic visual space was calculated by:

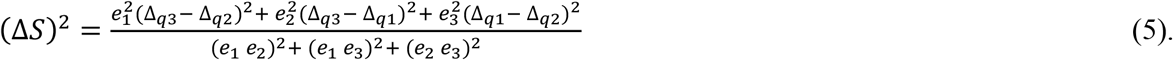

Grey distractor spectra (N=13) were chosen to be <1 Δ*S* of the achromatic point of *A. ocellaris* and ranged between 0.3 Δ*S* to 0.8 Δ*S* of each other. To control for the potential use of achromatic (intensity) cues when discriminating targets, we selected 6 to 10 distractor greys (from the 13) per stimulus based on all four-cone quantum catches to encompass the highest and lowest target intensities (see Supplementary Data 2 for distractor quantum catches and LED input values).

We chose target colors that increased in Δ*S* away from grey distractors in nine different directions within *A. ocellaris* color space. Each color set comprised of between 6 to 11 colors that varied from a high-saturated target color which was deemed highly contrasting against the grey distractors, to a low-saturated target color which had low contrast (<1 Δ*S*) against the grey distractors. The number of contrasts per color set included: ten for UV; six for UV-blue and purple; nine for blue, green, red, and violet-green; 11 for UV-red; and seven for orange. Note, we refer to colors as either non-UV colors (e.g., blue, green, red, purple, orange) or UV-colors (e.g., UV-blue, violet-green, UV-red) based on whether contrast was induced predominantly by UV/violet LED input. Color sets were plotted as lines in the receptor noise corrected space of *A. ocellaris* using σ = 0.11 for single cones and σ = 0.14 for double cones. To visualize the directionality of any asymmetrical contours in receptor space, an ellipsoid was manually created using Adobe Illustrator (2020) to be approximately centered on the average grey distractor point and intersect with the color discrimination thresholds.

Alternative models calculated Δ*S* values using more-conservative receptor σ-values ranging from 0.05 to 0.15, to assess their fit with *A. ocellaris* behavioral thresholds. Lower single cone noise (σ = 0.04 – 0.11) than double cones (σ = 0.14) was also modelled in case of different inherent noise levels. Threshold predictions were also compared between models of trichromat and tetrachromat vision in *A. ocellaris*, in case this could reveal any information on the contribution of double cones to color vision. The closest model fit was determined based on which had the smallest mean difference summed across all color lines from 1 Δ*S*.

Note that it is likely that color thresholds are not set by noise in individual photoreceptors but depend upon pooling of receptor signals across a fixed area of the retina (*39*). Consequently, thresholds should depend upon the density of a given receptor type in the retinal cone array (*39, 40*). If relative receptor densities vary across the retina [e.g., (*47, 77*)] this might lead to variations in color thresholds across the visual field.

### Calculating hue angles

To gain additional information on the perceptual properties of colors including the type of UV chromatic contrast and the presence of any complementary pairs, we calculated the elevation and azimuth angles of vectors plotted in anemonefish color space which corresponded to the psychophysical thresholds of color sets.

First, we converted ΔS to noise-corrected *XYZ* Cartesian coordinates using the ‘jnd2xyz’ function in the R package PAVO2 (85), which performs calculations based on the algorithm from (Pike, 2012; Maia & White, 2018). This returned XYZ coordinates for color threshold vectors representing the difference in receptor signal for the *x* axis [L – (M_1_ + M_2_)], *y* axis [M_1_ – (M_2_ + L)], and *z* axis [U – (M_1_ + M_2_ + L)].

We then calculated elevation angle ‘Θ’ by

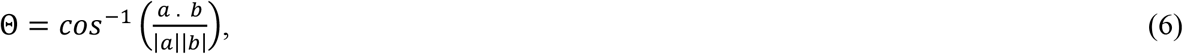

where ‘*α*. *b*’ was the dot product of vectors ‘*α*’ threshold *XYZ* position, and ‘*b*’ threshold *XYZ* position with the *Z* axis normal to the *XY* plane (i.e., zero), and |*α*| |*b*| were the magnitudes of each vector e.g.,

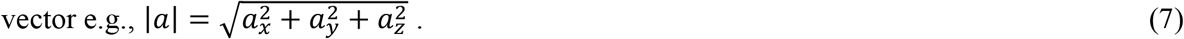

The product in units of radians was then converted to degrees and given appropriate signage to indicate relative position above (positive) or below (negative) the *XY* plane (−90° ≤ Θ ≤ 90°).

Thus, giving an elevation angle where the vertex was at the grey point (origin) and position was relative to an (*XY*) equator to indicate UV receptor stimulation based on movement along the *x* axis.

The azimuth angle ‘φ’ was calculated by

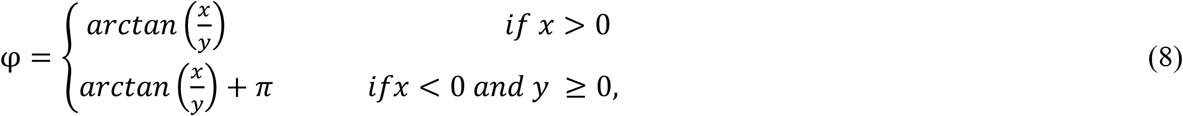

negative azimuth angles were corrected by adding 360°, so that all values ranged between 0° – 360°, where a bearing of 0°/360° marked the *x* axis describing L cone stimulation.

### Training and experiment

During both training and the experiment, the LED display was presented in a section of the aquarium separated by a sliding, opaque door. This door was closed to kept fish from viewing the display while the stimulus was updated between trials, and only upon trial commencement was the door raised to allow fish to view and interact with the display. For both training and testing, a morning (09:00 – 11:00) and afternoon (14:00 – 16:00) session were run, in which fish completed between 10 to 12 trials per day.

Fish were initially enticed to peck the LED display by presenting a pseudo-randomly chosen high contrast pixel (blue, green, red, or UV) with a small piece of prawn meat smeared on it. Over a week, we gradually reduced the size of the smeared food and transitioned towards a food-reward (Formula One Ocean Nutrition pellets) delivered by forceps when fish pecked the single target pixel. Once anemonefish readily approached and pecked at the display without enticement, we introduced the grey distractor pixels alongside the target pixel. Fish were only rewarded when they correctly chose/pecked the target color within 60 seconds. They were deemed to have reached the training criteria for the discrimination task after maintaining a correct choice probability of 0.75 over five consecutive sessions. 11 anemonefish met this criteria (mean number of training trials ± sd = 8.0 ± 4) and underwent experimental testing.

For testing, like training, fish were only rewarded for pecking the target pixel. Trials were terminated if fish made more than one incorrect choice or exceeded 60 seconds, upon which fish were returned to behind the divider (starting position) without reward. Note, because of the numerosity of pixels (n=38) per stimulus and the potential for distractions, each fish was permitted to make up to one incorrect choice per trial. For each trial, we recorded whether fish made a correct or incorrect choice, time (seconds) after fish entered through the door till target detection (i.e., latency), tested color set, and target Δ*S*.

Each color set was tested using five or six individual anemonefish that completed a minimum of eight trials per target color per assigned set (mean ± sd = 10 ± 1.0). Fish were divided into two groups assigned different color sets, including: *1*) Fish IDs 19, 20, 33, 34, and 36 which were assessed in order of testing with green, UV, purple, and UV-red, and *2*) Fish IDs 21, 22, 24, 31, 32, and 35 which were assessed in order of testing with blue, UV-blue, violet-green, red, and orange.

Between each trial the target pixel contrast was pseudo-randomly assigned from a list of LED intensity values for each color set. Throughout the experiment, we included control trials (n=10) to ensure that no other cues were created by the controller or code when choosing the target pixel, this determined the random chance of fish making a correct choice by displaying a target pixel of zero contrast (i.e., grey). In none of the control trials did fish correctly peck the control target.

To verify that differences in discrimination thresholds were not influenced by the order in which each of the color sets were tested, we reassessed each of the nine sets at the end of the experiment using two anemonefish from each group. Behavioral thresholds and psychometric functions from this secondary assessment were then compared with the primary assessment. Although we found evidence that experience effects had influenced the shape or incline of the psychometric function for some color sets (e.g., UV, blue, green, and UV-blue), there was none indicating that experience had contributed to differences in color discrimination thresholds that remained unchanged in the reassessment. The direction and size of differences among color discrimination thresholds did not vary systematically over the course of the study.

### Software and statistical analyses

All statistical analyses and color modelling were conducted using the statistical program R (v. 4.0.2) (*78*). Color distances were calculated using the RNL model and plotted in a tetrahedral space using the package ‘PAVO 2’ (*79*). Discrimination thresholds were determined by the point at which fish had a 0.5 probability of making a correct choice approximately at the inflection or steepest point of a sigmoid curve fitted to the behavioral data (Supplementary Data 4) using the package ‘quickpsy’ (*80*). A binomial test to determine the minimum number of correct choices required to be significantly above random choice (*48, 81*) gave a minimum threshold of discrimination at 20% of correct choices for our experiment (probability of one trial success =0.03, n = 10 trials per target, *p* = 0.04), indicating that our threshold criterion was conservative.

The effect of color sets on anemonefish discrimination thresholds was assessed using a linear mixed-effects model (LMM) run using function ‘lmer’ in the package ‘lme4’ (*82*). Individual threshold Δ*S* value was treated as the response variable, color sets as the fixed factor, and fish ID was the random effect. A post-hoc, pair-wise analysis controlled for multiple comparisons of threshold Δ*S* values across all possible combinations of color sets using Bonferroni adjustment (p.adjust, R base package ‘stats’). Another set of LMMs tested the effect of color on the duration (latency) of trials, where trial latency (in seconds) was the response variable, color set, target Δ*S* and the first-order interaction between color set and target Δ*S* were fixed effects, and fish ID was a random effect. Nine models were run (one per color set), where the intercept was assigned to a specific color to enable pairwise comparisons with the others.

To test whether the color set influenced the relationship between Δ*S* and the proportion of correct choices (i.e., did color influence how fish responded to the test and resulting shape of the psychometric curve), a generalised linear mixed effects model (GLMM) was run (*82*). Variables included anemonefish choice (0 = incorrect, 1 = correct) used as a binomial response variable, Δ*S*, color set and the first-order interaction between the two variables treated as fixed factors, and Fish ID entered as a random effect. Model p-values were corrected for multiple comparisons via Bonferroni adjustment (‘p.adjust’, base R package ‘Stats’).

Residual diagnostics were performed for LMMs and GLMM using the package ‘DHARMa’ (*83*), that checked the distribution of residuals and verified there were no dispersion issues.

Analysis of anemonefish skin reflectance and target color emission were performed using the ‘vismodel’ and ‘spec2rgb’ functions in the package ‘PAVO 2’ (*79*).

## Acknowledgments

We would like to thank Adélaιde Sibeaux for giving advice on the statistical analysis. We thank Aidan McGuire for assisting with the experiment. We also thank all volunteers for their aid in maintaining aquaria.

## Author contributions

Conceptualization: KLC, LJM, FC, NJM, AP

Methodology: KLC, LJM, FC, NJM, AP, WC

Investigation: LJM, AP, WC, NJM

Visualization: LJM, KLC, DCO

Supervision: KLC

Writing—original draft: LJM

Writing—review & editing: LJM, KLC, DCO, FC, NJM, WC, AP

## Data and materials availability

All supporting data are available in the supplementary materials, and raw data can be accessed from The University of Queensland’s eSpace data repository via (TBA).

