## Supplementary for "Anemonefish have finer color discrimination in the ultraviolet"

Laurie J Mitchell *et al.*

**This PDF file includes:**

Figs. S1 to S6
Tables S1

**Other Supplementary Materials for this manuscript include the following:**

Data S1 to S3

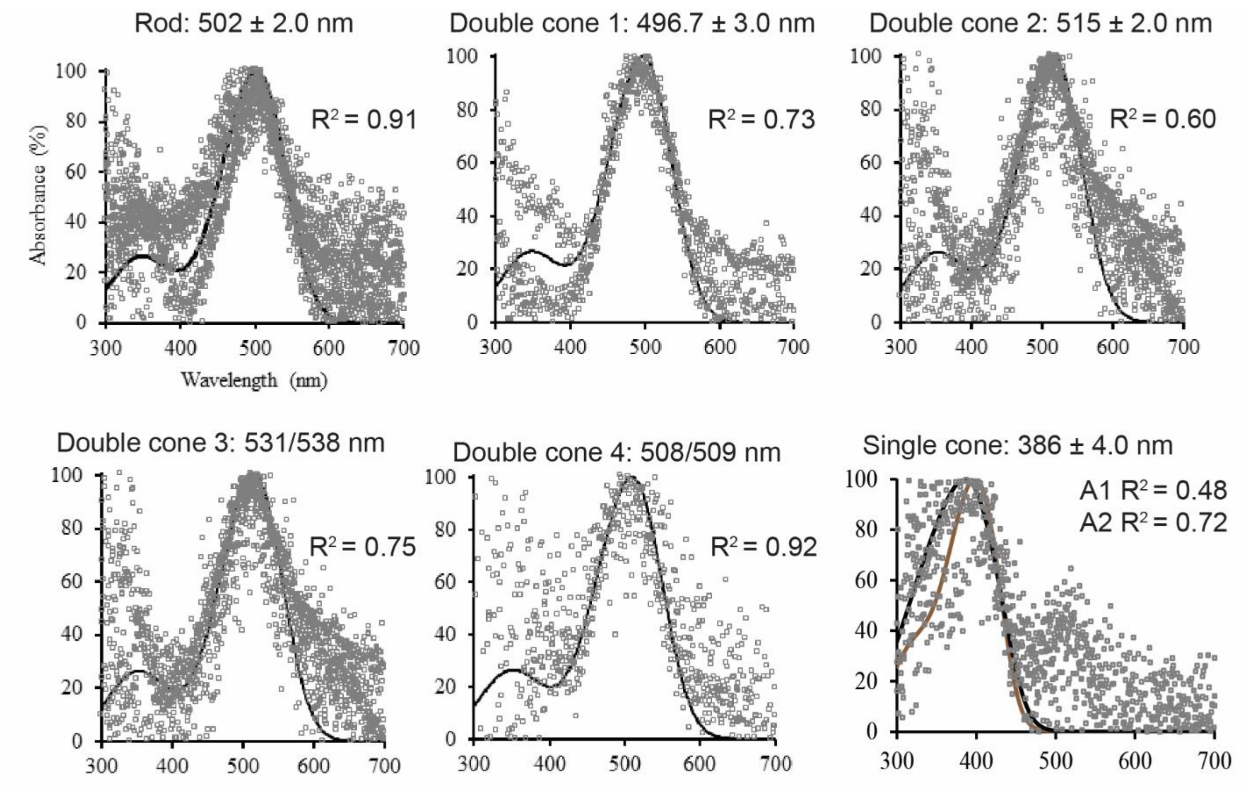


**Fig. S1.**

**Photoreceptor spectral absorbance measurements from miscrospectrophotometry.**

Normalised individual absorbance spectra from MSP including rods (n=7), double cones (1, n = 4; 2, n = 6; 3, n = 2; and 4, n = 2), and single cones (n=4). Given are average $\lambda_{max}$ values ± s.e.m. R-squared values were calculated by least squares regression using averaged absorbance spectra. Note: each subplot contains a fitted A1 visual pigment template, except for the single cone which has both fitted A1 (brown line curve) and A2 (black line curve) visual pigment templates for comparison. Individual absorbance measurements and lens transmittance data can be accessed via S3 Data.

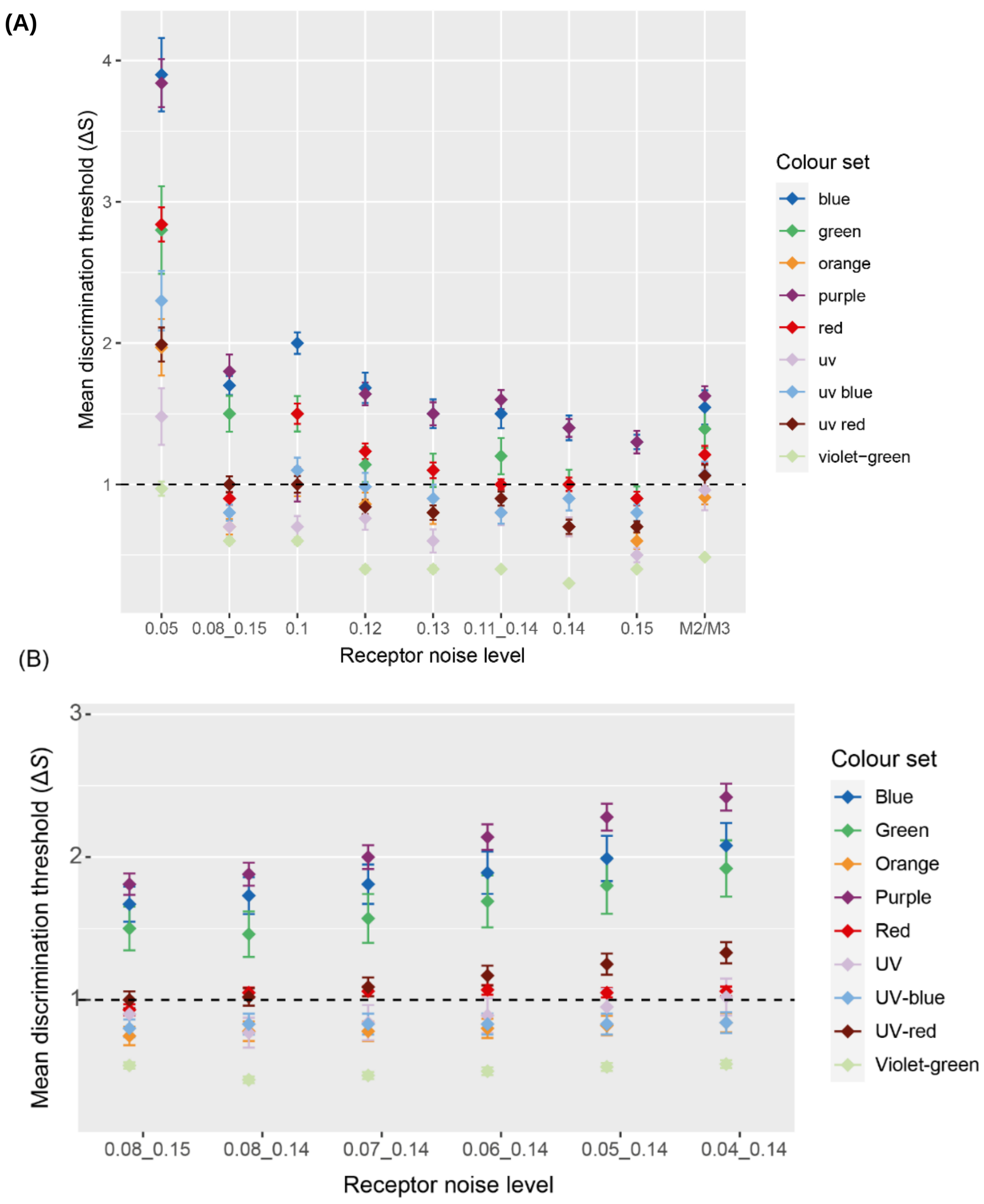


**Fig. S2.**

**RNL predicted color discrimination thresholds using different receptor noise levels. (A)**

Comparison of colour discrimination thresholds ($\Delta S$) calculated using a range of receptor noise ($\sigma$) levels, with (or without) input by different MWS cones (using $\sigma$ value = 0.14), and **(B)** additional distinct receptor noise levels for single cones and double cones. Initial and secondary receptor noise values (‘X.XX_X.XX’) refer to single cones and double cones, respectively. Discrimination thresholds were averaged across anemonefish for the $\Delta S$ which corresponded to a

50% proportion of correct choices. Error bars are the s.e.m. Broken, horizontal line demarks the RNL model assumed threshold ($\Delta S = 1$). ‘M2/M3’ refers to modelling with the $M_3$ cone in place of the $M_2$ cone.

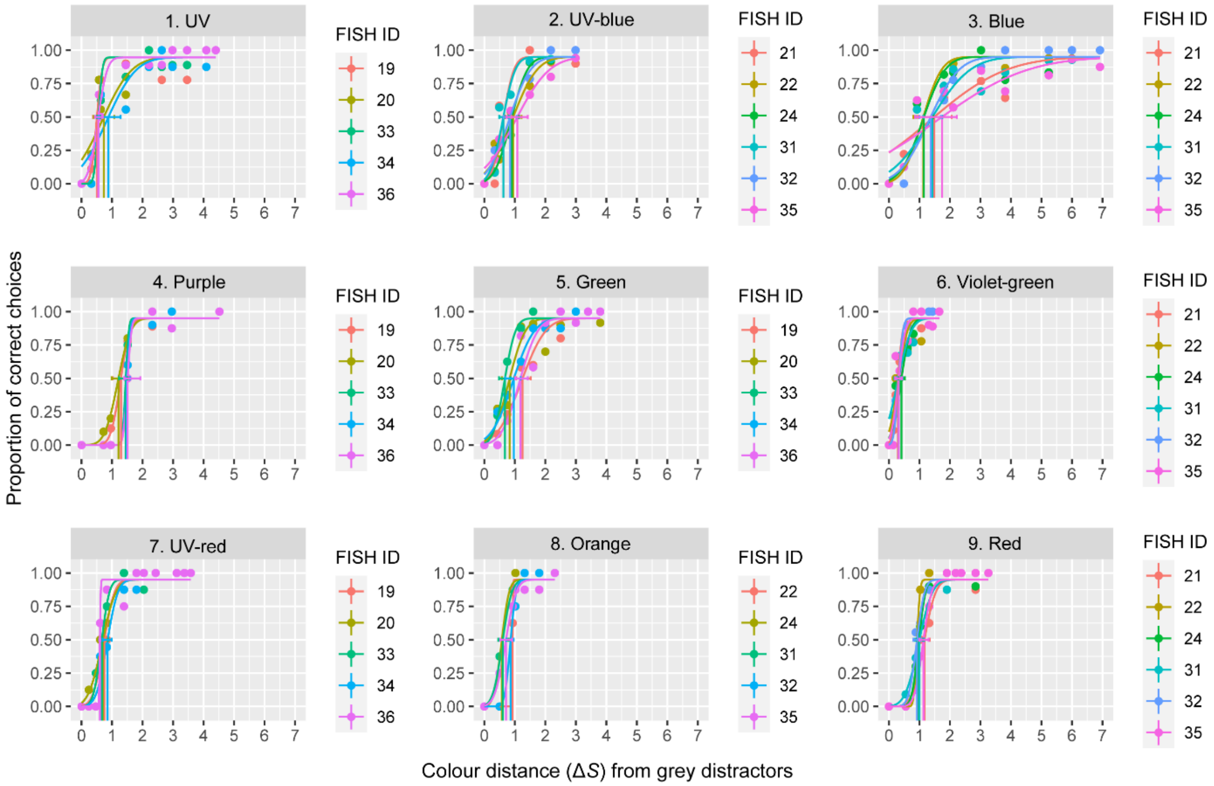


**Fig. S3.**

**Non-truncated individual discrimination thresholds**. Color discrimination thresholds shown as a function of the proportion of correct choices by anemonefish for targets with a range of chromatic contrasts (ΔS). Error bars denote 0.95 CIs. Discrimination thresholds are values calculated per fish (Fish ID) and are demarked by vertical lines. Each plotted point represents the mean proportion of correct choices from one fish (n = 8 to 17 trials).

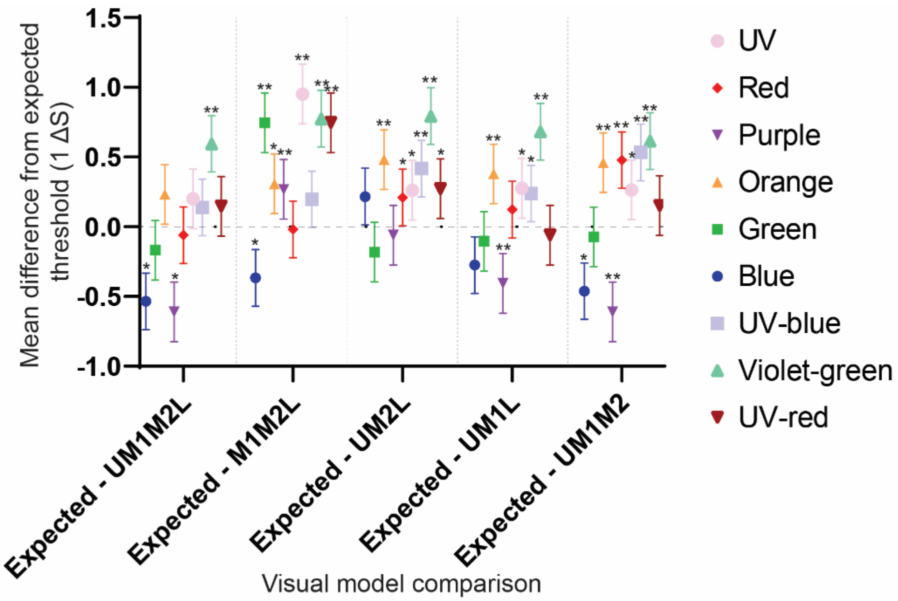


| Colour set | RNL Model | Mean | Difference | SE of difference | t ratio | df | P value | Adjusted P Value |
|---|---|---|---|---|---|---|---|---|
| Blue | UM1M2L | 1.535 | 0.535 | 0.1033 | 5.179 | 5 | 0.003529 | 0.031757 |
| | M1M2L | 1.367 | 0.3667 | 0.06075 | 6.035 | 10 | 0.000126 | 0.001135 |
| | UM2L | 0.7833 | -0.2167 | 0.07923 | 2.735 | 10 | 0.021028 | 0.18925 |
| | UM1L | 1.275 | 0.275 | 0.09298 | 2.958 | 10 | 0.014345 | 0.129106 |
| | UM1M2 | 1.462 | 0.4617 | 0.1134 | 4.072 | 10 | 0.002244 | 0.020197 |
| Green | UM1M2L | 1.168 | 0.168 | 0.1282 | 1.311 | 4 | 0.260085 | >0.999999 |
| | M1M2L | 0.254 | -0.746 | 0.01796 | 41.53 | 9 | <0.000001 | <0.000001 |
| | UM2L | 1.182 | 0.182 | 0.1143 | 1.593 | 9 | 0.145673 | >0.999999 |
| | UM1L | 1.104 | 0.104 | 0.1029 | 1.01 | 9 | 0.33876 | >0.999999 |
| | UM1M2 | 1.072 | 0.072 | 0.1131 | 0.6365 | 9 | 0.540299 | >0.999999 |
| Orange | UM1M2L | 0.768 | -0.232 | 0.07864 | 2.95 | 4 | 0.041961 | 0.377645 |
| | M1M2L | 0.692 | -0.308 | 0.06983 | 4.411 | 9 | 0.001693 | 0.015241 |
| | UM2L | 0.518 | -0.482 | 0.04 | 12.05 | 9 | <0.000001 | 0.000007 |
| | UM1L | 0.62 | -0.38 | 0.05263 | 7.22 | 9 | 0.00005 | 0.000448 |
| | UM1M2 | 0.54 | -0.46 | 0.02211 | 20.8 | 9 | <0.000001 | <0.000001 |
| Purple | UM1M2L | 1.61 | 0.61 | 0.06753 | 9.033 | 4 | 0.000832 | 0.007487 |
| | M1M2L | 0.732 | -0.268 | 0.03053 | 8.778 | 9 | 0.00001 | 0.000094 |
| | UM2L | 1.06 | 0.06 | 0.03473 | 1.728 | 9 | 0.118094 | >0.999999 |
| | UM1L | 1.406 | 0.406 | 0.05503 | 7.378 | 9 | 0.000042 | 0.000378 |
| | UM1M2 | 1.61 | 0.61 | 0.06096 | 10.01 | 9 | 0.000004 | 0.000032 |
| Red | UM1M2L | 1.06 | 0.06 | 0.03578 | 1.677 | 5 | 0.154377 | >0.999999 |
| | M1M2L | 1.018 | 0.01833 | 0.04833 | 0.3793 | 10 | 0.712388 | >0.999999 |
| | UM2L | 0.79 | -0.21 | 0.03786 | 5.547 | 10 | 0.000245 | 0.002207 |
| | UM1L | 0.8767 | -0.1233 | 0.03947 | 3.125 | 10 | 0.010785 | 0.097063 |
| | UM1M2 | 0.5217 | -0.4783 | 0.006009 | 79.6 | 10 | <0.000001 | <0.000001 |
| UV | UM1M2L | 0.8 | -0.2 | 0.08944 | 2.236 | 4 | 0.089009 | 0.801084 |
| | M1M2L | 0.0488 | -0.9512 | 0.0008752 | 1087 | 9 | <0.000001 | <0.000001 |
| | UM2L | 0.738 | -0.262 | 0.06977 | 3.755 | 9 | 0.004518 | 0.040661 |
| | UM1L | 0.724 | -0.276 | 0.07202 | 3.832 | 9 | 0.004014 | 0.036128 |
| | UM1M2 | 0.736 | -0.264 | 0.06931 | 3.809 | 9 | 0.00416 | 0.037443 |
| UV-blue | UM1M2L | 0.8617 | -0.1383 | 0.07897 | 1.752 | 5 | 0.140207 | >0.999999 |
| | M1M2L | 0.8033 | -0.1967 | 0.07032 | 2.797 | 10 | 0.018896 | 0.170062 |
| | UM2L | 0.5833 | -0.4167 | 0.03073 | 13.56 | 10 | <0.000001 | <0.000001 |
| | UM1L | 0.7617 | -0.2383 | 0.0621 | 3.838 | 10 | 0.003276 | 0.02948 |
| | UM1M2 | 0.4667 | -0.5333 | 0.04425 | 12.05 | 10 | <0.000001 | 0.000003 |
| Violet-green | UM1M2L | 0.405 | -0.595 | 0.02217 | 26.83 | 5 | 0.000001 | 0.000012 |
| | M1M2L | 0.225 | -0.775 | 0.01544 | 50.2 | 10 | <0.000001 | <0.000001 |
| | UM2L | 0.2067 | -0.7933 | 0.006667 | 119 | 10 | <0.000001 | <0.000001 |
| | UM1L | 0.3183 | -0.6817 | 0.01641 | 41.53 | 10 | <0.000001 | <0.000001 |
| | UM1M2 | 0.385 | -0.615 | 0.0263 | 23.38 | 10 | <0.000001 | <0.000001 |
| UV-red | UM1M2L | 0.854 | -0.146 | 0.05056 | 2.888 | 4 | 0.04466 | 0.401939 |
| | M1M2L | 0.254 | -0.746 | 0.01796 | 41.53 | 9 | <0.000001 | <0.000001 |
| | UM2L | 0.728 | -0.272 | 0.04407 | 6.171 | 9 | 0.000164 | 0.00148 |
| | UM1L | 1.06 | 0.06 | 0.06122 | 0.98 | 9 | 0.352676 | >0.999999 |
| | UM1M2 | 0.848 | -0.152 | 0.04295 | 3.539 | 9 | 0.006324 | 0.05692 |

**Fig. S4.**

**Fit comparison of RNL predicted discrimination thresholds using tetrachromatic and trichromatic models**. Individual color set comparisons for tetrachromatic (UM1M2L) and trichromatic modelling in the accuracy of predicting color discrimination thresholds. For each model, the individual thresholds ($\Delta S$) per color set were subtracted from the expected threshold value of 1 $\Delta S$ (demarked by the broken line), and the overall average difference was calculated. Visual models either included input from all cones (tetrachromatic) or dropped input from one of the four cone types (trichromatic): UV-sensitive cone ‘U’, medium-wavelength sensitive cone 1 ‘M1’, medium-wavelength sensitive cone 2 ‘M2’, or long-wavelength sensitive cone ‘L’. Error bars indicate upper and lower 0.95 confidence limits. ‘ns’ indicates $p > 0.05$, ‘*’ indicates $p = 0.05 > 0.001$, and ‘**’ indicates $p < 0.001$. The summary table contains the results of a two-way ANOVA with Dunnett’s multiple comparisons test.

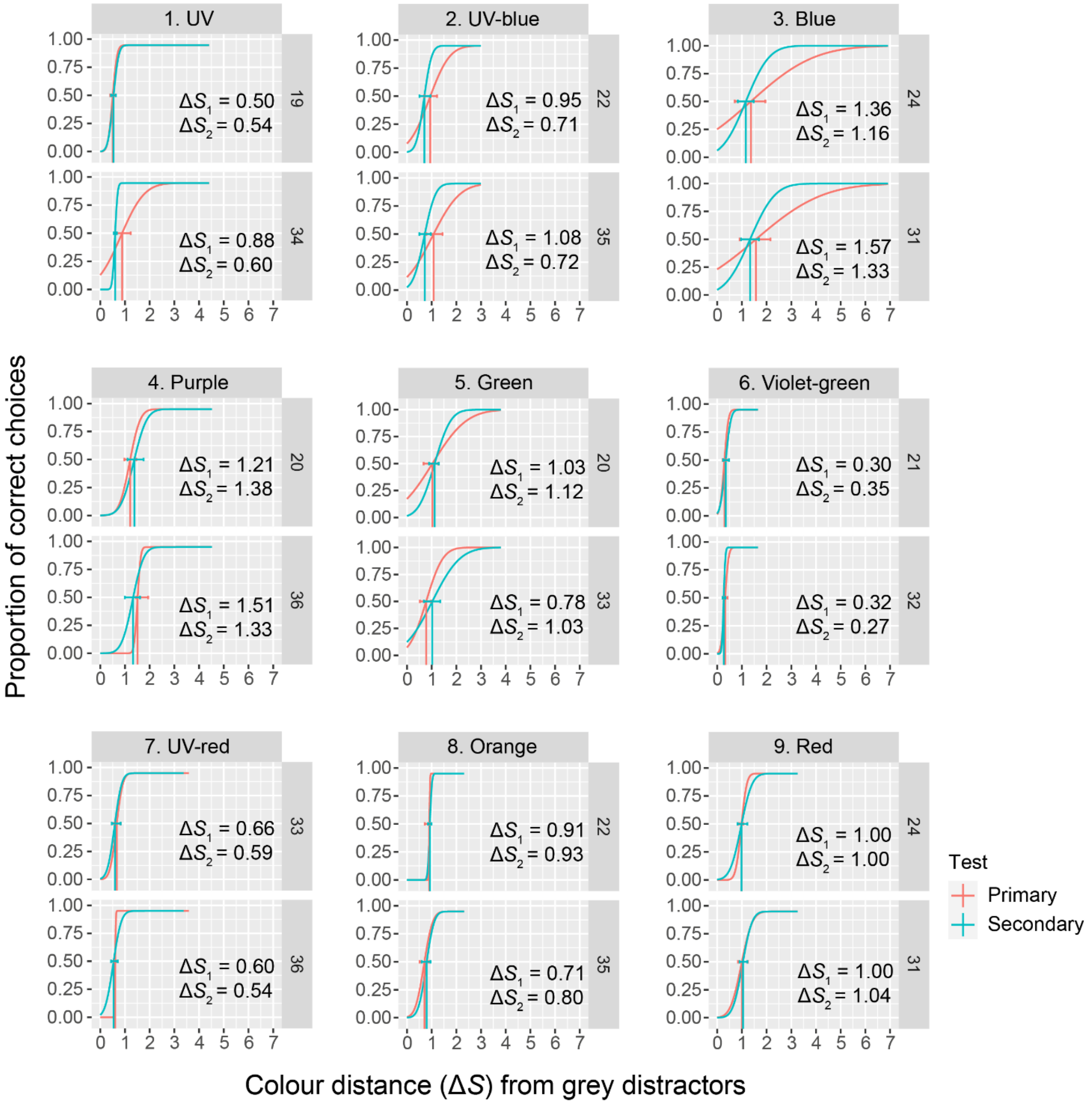


**Fig. S5.**

**Discrimination thresholds compared between primary and secondary assessment trials.**

Comparisons between anemonefish (n = 2) discrimination thresholds for primary (ΔS1) and secondary (ΔS2) assessments of the nine color sets to assess for the effect of experience on performance. Psychometric curves in each subplot depict the proportion of correct choices by individual anemonefish (fish ID labelled on the right-hand side) as a function of ΔS per color set. Vertical lines demark the target ΔS which corresponded to a 0.5 chance of making a correct choice. Color distances were calculated using the RNL model with a σ value of 0.14.

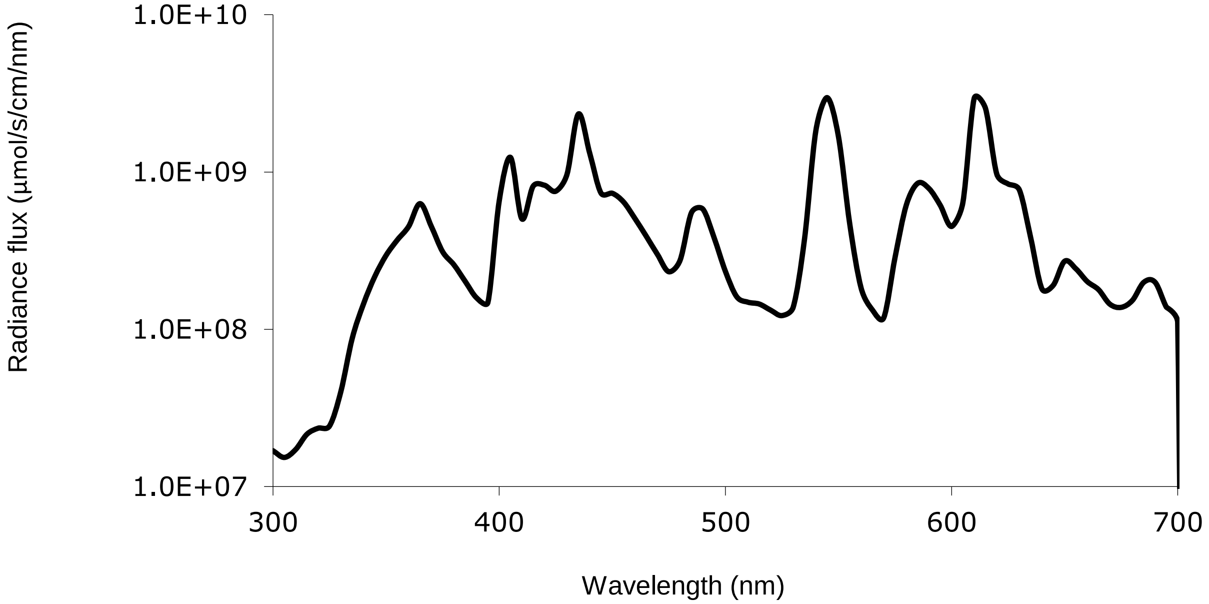


**Fig. S6**

**Experimental background illumination.** Side-welling measurement of the radiance flux (photons/s/cm/nm) of the PTFE LED display screen/background under the aquarium lighting in which colors were presented. Note, y-axis units are in log-scale due to the sharp order of magnitude fluctuations of the fluorescent light that otherwise obscures the remainder of the spectrum.

**Table S1.**

**Relative cone quantum catches and XYZ coordinates for threshold loci.** Relative quantum catches ($q_i$) for cones and cartesian XYZ vector coordinates for the threshold ΔS of each colour set and the averaged grey distractor.

| Colour set | $q_u$ | $q_{m1}$ | $q_{m2}$ | $q_l$ | X | Y | Z |
|---|---|---|---|---|---|---|---|
| Blue | 0.238458 | 0.266192 | 0.257159 | 0.238191 | -0.63157 | 0.876254 | -0.77094 |
| Green | 0.228171 | 0.242855 | 0.263887 | 0.265088 | 0.259796 | 0.080515 | -1.02514 |
| Orange | 0.257067 | 0.222486 | 0.240732 | 0.279716 | 0.400307 | -0.72128 | 0.271744 |
| Purple | 0.225555 | 0.271354 | 0.257999 | 0.245092 | -0.70295 | 0.768823 | -1.33153 |
| Red | 0.250623 | 0.219724 | 0.237959 | 0.291694 | 0.527345 | -1.1117 | 0.08149 |
| UV | 0.278353 | 0.235382 | 0.244059 | 0.242207 | -0.05871 | 0.267704 | 0.943686 |
| UV-blue | 0.264361 | 0.253696 | 0.248361 | 0.233581 | -0.5153 | 0.746421 | 0.307825 |
| Violet-green | 0.265061 | 0.231425 | 0.251161 | 0.252353 | 0.258383 | 0.075514 | 0.49614 |
| UV-red | 0.275431 | 0.229926 | 0.242762 | 0.251881 | 0.124372 | -0.04697 | 0.884096 |
| Average grey | 0.257962 | 0.239286 | 0.251447 | 0.251305 | 0.036433 | 0.193739 | 0.168948 |

Cone types are ultraviolet-sensitive ‘u’, medium-wavelength sensitive 1 ‘$m_1$’ and 2 ‘$m_2$’, and long-wavelength sensitive ‘l’.

**Data S1. (separate file)**

Results from linear mixed effects models (LMM) and generalized linear mixed effects model (GLMM).

**Data S2. (separate file)**

Calculated absolute quantum catches for anemonefish cones in elicited by the LED emission of target colors and distractor greys.

**Data S3. (separate file)**

Averaged and fitted spectral absorbance data for *A. ocellaris* cones, including individual spectral absorbance measurements from microspectrophotometry and lens transmission measurements.

**Data S4. (separate file)**

Behavioral dataset used in the analysis of the color discrimination experiment.
